# Important Docking study of Structure of Plasmodium falciparum thioredoxin reductase-thioredoxin complex: the case of potent drug Telatinib, a small molecule angiogenesis inhibitor

**DOI:** 10.1101/2021.07.24.453649

**Authors:** Ivan Vito Ferrari

## Abstract

**Background:** Over the last decades, malaria parasites have been rapidly developing resistance against antimalarial drugs, which underlines the need for novel drug targets. Thioredoxin reductase (TrxR) is crucially involved in redox homeostasis and essential for Plasmodium falciparum. In this communication, we report first time important Docking study by in Silico approach, using AutoDock Vina. After a selective analysis of over 300 drugs, processed with Pyrx (a Virtual Screening software into the active site of protein (ID PDB 4J56 Thioredoxin reductase 2 Chain A), we noticed excellent value of Binding Energy of Telatinib estimated by Pyrx software. These results are comparable to the crystallized ligand FAD (FLAVIN-ADENINE DINUCLEOTIDE) completed in the above-mentioned protein. Indeed, from the results of Autodock Vina, Telatinib an inhibitor of tyrosine kinases, has excellent a Binding affinity value, ca. −12 kcal/mol.

## 1. Introduction

Over the last decades, malaria parasites have been rapidly developing resistance against antimalarial drugs, which underlines the need for novel drug targets. Thioredoxin reductase (TrxR) is crucially involved in redox homeostasis and essential for Plasmodium falciparum[1]. Plasmodium falciparum is a unicellular protozoan parasite of humans, and the deadliest species of Plasmodium that causes malaria in humans. The parasite is transmitted through the bite of a female Anopheles mosquito and causes the disease’s most dangerous form, falciparum malaria [2]. It is responsible for around 50% of all malaria cases. Plasmodium falciparum is the Plasmodium species responsible for 85 % of the malaria cases. Malaria infects over 200 million people annually, mostly in poor tropical and subtropical countries of Africa [2–3]. In this short communication, we investigated about 300 drugs, through In Silico Docking approach, downloaded from PubChem Database (https://pubchem.ncbi.nlm.nih.gov/). We have focused on Autodock Vina, estimated with Pyrx software, a simple Virtual Screening library software for Computational Drug Discovery (https://pyrx.sourceforge.io/), based on prediction the binding orientation and affinity of a ligand [4].

## 2. Materials and methods

### 2.1 Protein and Ligand Preparation before docking

#### Protein Preparation

**4J56** [Structure of Plasmodium falciparum thioredoxin reductase-thioredoxin complex: Thioredoxin reductase 2] was prepared manually using several software, before molecular docking analysis. The first step, was downloaded from Protein Data Bank, https://www.rcsb.org/structure/4j56 and save in pdb format. The second step were removed all unnecessary docking chains. In fact, in this case we are only Chain A (https://www.rcsb.org/structure/6LD3) has been maintained and re-saved in pdb format. (See below figure 1) Next, were the removal of ligands and water molecules crystallized using Chimera software [6]. Later, polar hydrogens and Kollmann charges were added with Mgl-Tool, (or called AutoDockTools, a software developed at the Molecular Graphics Laboratory (MGL) [7] As a last step they were added to the protein, any missing amino acids and the whole protein was minimized with the Swiss PDB Viewer Software [8].

**Fig 1.**
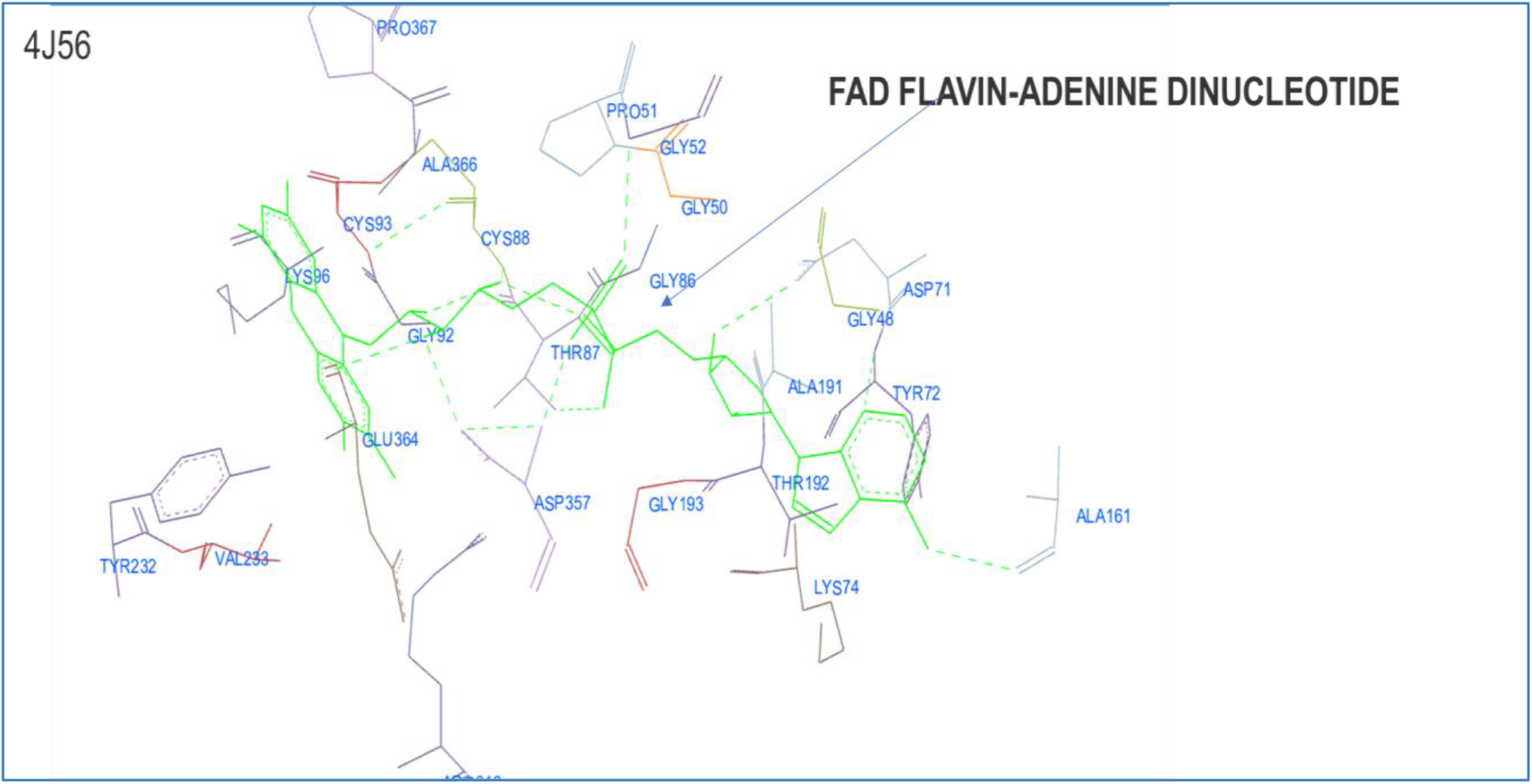
3D Structure of ID PDB 4J56 Chain A (Structure of Plasmodium falciparum thioredoxin reductase-thioredoxin complex by Discovery Studio Biovia Software [10]

##### Ligand Preparation

the first step, was to separate the crystallized ligand G8O (2,4-dimethoxy-5-thiophen-2-yl-benzoic acid) from its protein, manually add all the hydrogens and their Gasteiger charges (with the MGL Tool software) and minimize it with MMFF94 force field [9], and opened by Pyrx software (https://pyrx.sourceforge.io/).

Parameters Grid Box for Docking in Ligand Binding Site Pocket for Repeatability Binding Affinity by AMDock Software calculated with Autodock Vina and Autodock 4:

**- ID PDB 4J56: Structure of Plasmodium falciparum thioredoxin reductase-thioredoxin complex Chain A**: Center X (= −32.19); Centre Y (=-109.21); Centre Z (=196.34); Dimensions (Angstrom) (Å) X, Y, Z [= 26.70=,25.00, =20.53]; exhaustiveness = 8.

## 3 Discussion and Results

In this communication, we investigated about 300 drugs, through In Silico Docking approach, downloaded from PubChem Database (https://pubchem.ncbi.nlm.nih.gov/). From our results of Autodock Vina, estimated with Pyrx software, a simple Virtual Screening library software for Computational Drug Discovery (https://pyrx.sourceforge.io/), based on prediction the binding orientation and affinity of a ligand [4] (**See below table 1**) we have selected only best Binding Energy of drugs (kcal/mol) (**See below table 2**). After intensive docking investigation by Autodock Vina, we propose a potential candidate drug against Malaria, that is Telatinib, C_20_H_16_ClN_5_O_3_. (**See below figure 2**) It is an orally administered drug that inhibits VEGFR-2, VEGFR-3 PDGFR-β and c-Kit (Eskens et al., 2009) and this drug is an orally available, potent multitargeted VEGFR-2, VEGFR-3, PDGFR-β and c-Kit tyrosine kinases inhibitor with an IC50 of 19 nM for the inhibition of VEGFR-2 autophosphorylation and it is under investigation in clinical trial NCT03817411 (Telatinib in Combination With Capecitabine/ Oxaliplatin in 1st Line Gastric or GEJ Cancer [11–12]. We noticed excellent value of Binding Energy (kcal/mol) of Telatinib by docking analysis. These results are are comparable to the crystallized ligand FAD (FLAVIN-ADENINE DINUCLEOTIDE) completed in the above-mentioned protein. Indeed, from the results of Autodock Vina, estimated by Pyrx, Telatinib has excellent Binding affinity value, ca. −12 kcal/mol. (**See below Table 2**).

**Table 1.**
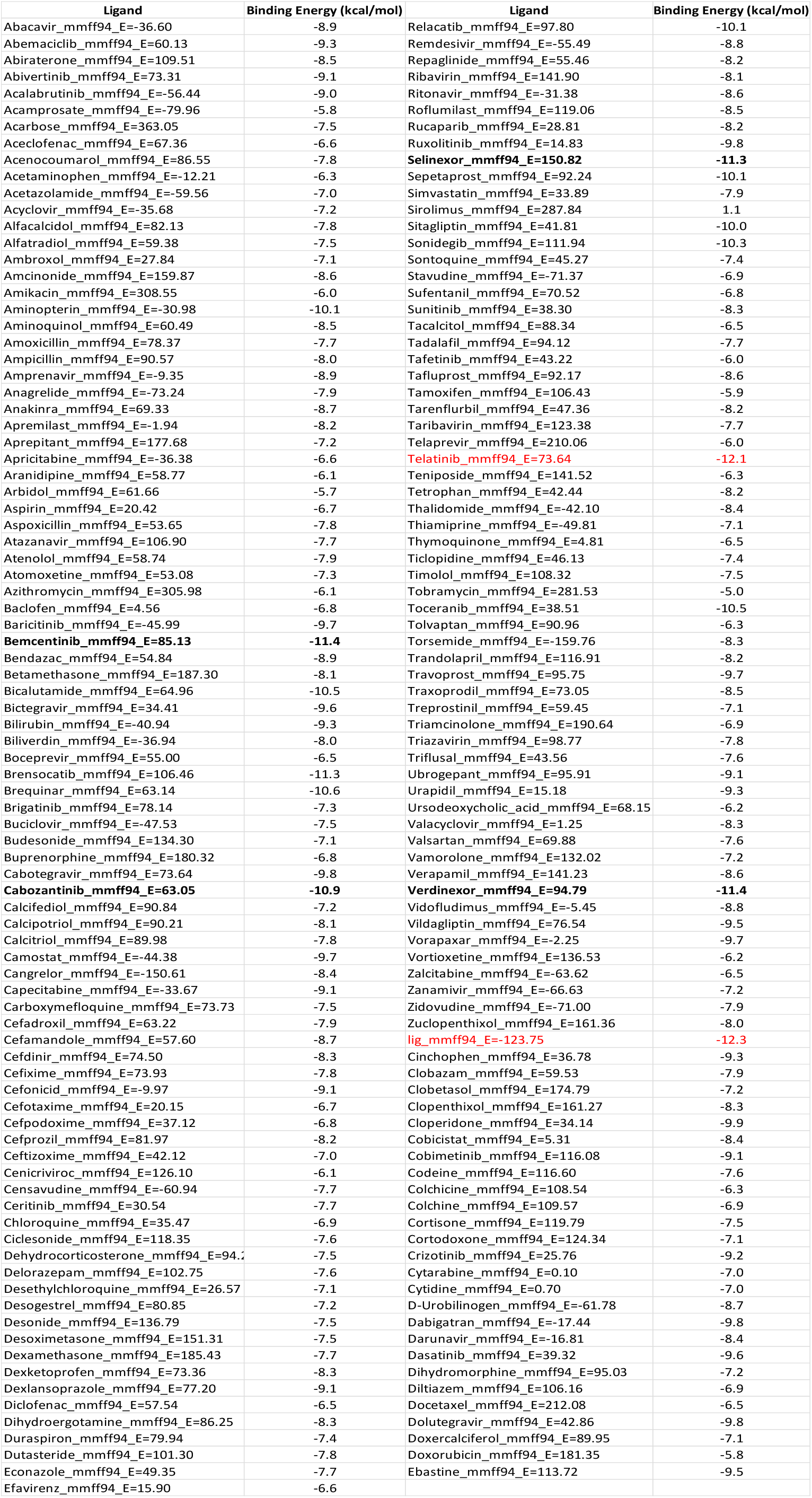
Binding energies (Kcal/mol) of Drugs into the active site of ID PDB 4J56 protein estimated by Autodock Vina with Pyrx software

**Table 2.**
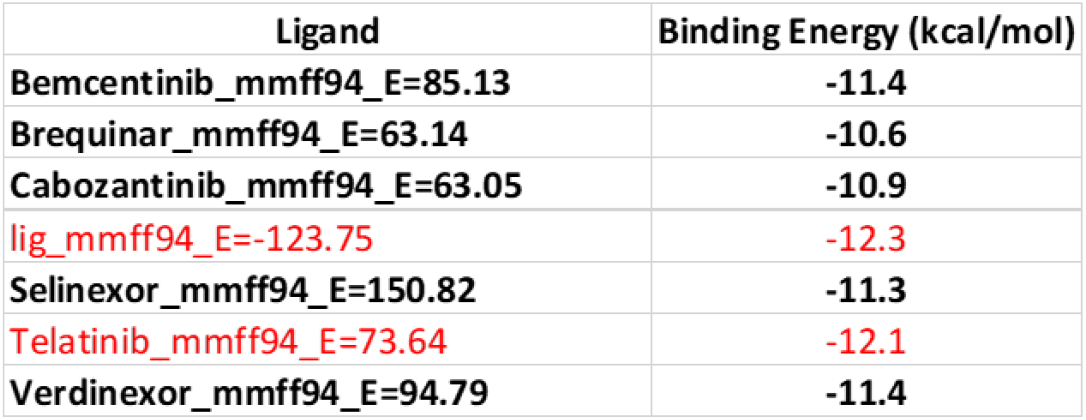
Best Binding energies (Kcal/mol) of Drugs into the active site of ID PDB 4J56 protein estimated by Autodock Vina with Pyrx software

| Ligand | Binding Energy (kcal/mol) |
| --- | --- |
| Bemcentinib_mmff94_E=85.13 | -11.4 |
| Brequinar_mmff94_E=63.14 | -10.6 |
| Cabozantinib_mmff94_E=63.05 | -10.9 |
| lig_mmff94_E=-123.75 | -12.3 |
| Selinexor_mmff94_E=150.82 | -11.3 |
| Telatinib_mmff94_E=73.64 | -12.1 |
| Verdinexor_mmff94_E=94.79 | -11.4 |

**Fig. 2.**
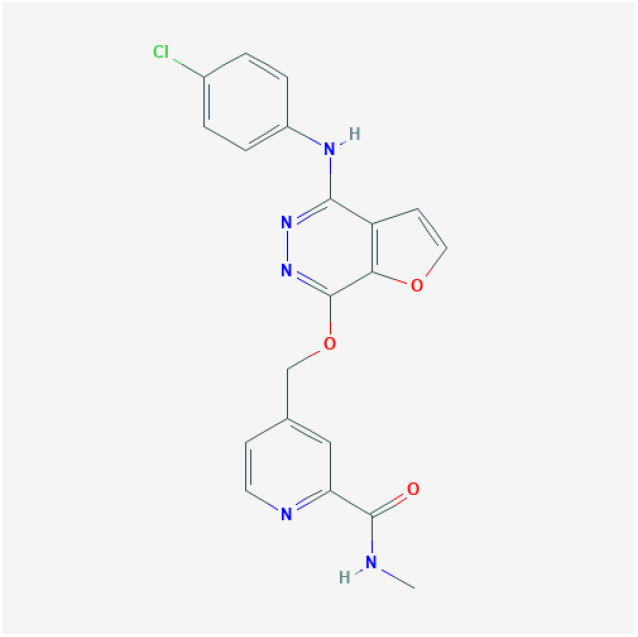
Structure of Telatinib

**Fig. 3.**
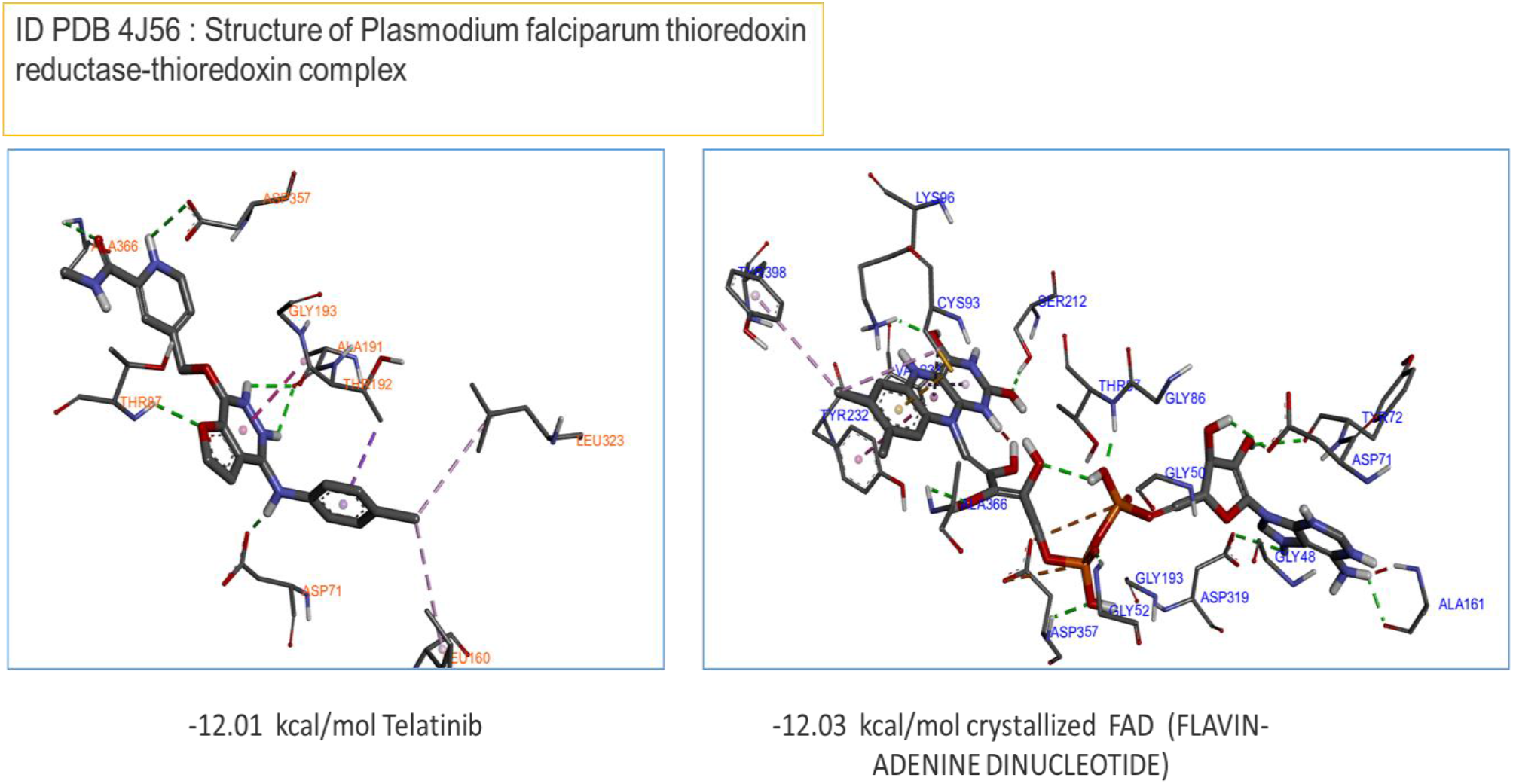
Comparison Binding Energy kcal/mol of Crystalized FAD – 12,0 kcal/mol with proposed drug Telatinib −12,01 kcal/ml into the active site of protein ID PDB: “Structure of Plasmodium falciparum thioredoxin reductase-thioredoxin complex”, calculated by Autodock Vina with Pyrx software. This figure was produced by Discovery Studio Biovia Software [10]

## 4. Conclusion

In this communication, we report first time important Docking study by in Silico approach, using AutoDock Vina. After a selective analysis of over 300 drugs, processed with Pyrx (a Virtual Screening software into the active site of protein (ID PDB 4J56 Thioredoxin reductase 2 Chain A), we noticed excellent value of Binding Energy of Telatinib estimated by Pyrx software. These results are comparable to the crystallized ligand FAD (FLAVIN-ADENINE DINUCLEOTIDE) completed in the above-mentioned protein. Indeed, from the results of Autodock Vina, Telatinib an inhibitor of tyrosine kinases, has excellent a Binding affinity value, ca. −12.01 kcal/mol versus −12.03 kcal/mol of crystallized ligand FAD.

## Author contributions

I.V.F. conceived, designed and wrote the paper, performed the calculations and analysed the data.

## Declaration of Competing Interest

The authors declare they have no potential conflicts of interest to disclose.

## Notes

### Competing Interest Statement

The authors have declared no competing interest.

